# Stochastic Gene Expression under Sequestration: Noise Reduction and Emergent Distributions

**DOI:** 10.1101/2025.11.24.690157

**Authors:** Olha Morozova, Ivana Oravcová, Iryna Zabaikina, Pavol Bokes, Abhyudai Singh

## Abstract

Gene expression noise can be modulated by protein sequestration, a mechanism we investigate through a stochastic modeling framework. We examine how the distribution of free (non-sequestered) protein depends on sequestration cooperativity (monomers, dimers, multimers) and on the timescale separation between sequestration and protein turnover. For non-cooperative sequestration, faster kinetics drive the distribution from a high-noise to a lower-noise gamma form, while the right-tail remains governed by the high-noise limit — revealing a non-commutativity between tail asymptotics and fast sequestration. For cooperative sequestration, the distribution departs from gamma, exhibiting left skewness or multi-modality. These results highlight how sequestration mechanisms shape protein variability in nontrivial ways.

## 1 Introduction

Randomness in biochemical reactions makes gene expression inherently stochastic, even in tightly controlled environments and clonal populations (Raj and Van Oudenaarden, 2008; Kaern et al., 2005; Elowitz et al., 2002; Singh and Hespanha, 2007). A major contributor to variability is bursting, wherein short episodes of high transcriptional/translational activity are interspersed with quiescent periods (Dar et al., 2012; Rodriguez and Larson, 2020; Chong et al., 2014; Suter et al., 2011; Fukaya, 2023; Ma et al., 2021; Brouwer et al., 2023; Raj et al., 2006; Fraser et al., 2021). Minimal bursty models—continuous piecewise deterministic processes or discrete birth–death processes—capture essential features of protein statistics, including gamma/negative-binomial stationary distributions that connect directly to burst frequency and size (Friedman et al., 2006; Paulsson and Ehrenberg, 2000; Ghusinga and Singh, 2015). Piecewise deterministic formulations can be derived from fully discrete models by scale separation arguments (Fang et al., 2020; Jia et al., 2019).

Beyond synthesis and dilution/decay, cells deploy *sequestration* to modulate protein activity: factors bind to decoy DNA sites, assemble into higher-order complexes, or compartmentalise into phase-separated condensates (Burger et al., 2012; Dey et al., 2020; Lee and Maheshri, 2012; Klosin et al., 2020; Buchler and Cross, 2009; Biswas et al., 2024). Such processes buffer fluctuations, reshape effective reaction rates, and can dynamically redistribute molecules between functionally active and inactive pools. These results motivate a systematic characterization of how cooperativity (monomers versus multimers) and kinetic timescales (fast versus slow sequestration relative to protein turnover) shape full steady-state distributions of the *free* (non-sequestered) protein.

Homeostasis is fundamental to cellular regulation and robust function. Perfect adaptation of the steady-state protein mean has been shown in antithetic integral feedback systems (Briat et al., 2016), and its impact on full protein distributions has been analyzed in (Briat et al., 2018). In the sequestration circuit, the free-protein mean is independent of sequestration kinetics, and we ask how cooperativity and time-scale reshape the distribution beyond its mean.

Timescale separation is particularly consequential. In classical bursty models without sequestration, deterministic decay between random bursts yields explicit gamma laws for protein concentration (Friedman et al., 2006). However, tail probabilities can be controlled by rare events whose asymptotics need not commute with fast-reaction limits, as shown for drift–jump systems subject to large deviations (Bokes, 2022).

Building on the above context, we develop a stochastic framework for gene expression under sequestration and report the following results:

### 1. Non-cooperative sequestration

As sequestration becomes fast relative to protein turnover, the free-protein distribution transitions from a high-noise regime to a lower-noise gamma law. Strikingly, the right tail remains governed by the high-noise limit, revealing a non-commutativity between fast-sequestration limits and tail large deviations. This tail divergence arises from the transient disruption of the sequestration quasi-steady state immediately after a burst event.

### 2. Cooperative sequestration

Dimeric and multimeric binding drive qualitative departures from gamma behavior, producing pronounced left-skewness or multimodality via nonlinear depletion of the active pool.

### 3. Design map for noise control

We chart parameter regimes, indexed by intrinsic noise and sequestration affinity, that either reduce noise under sequestration or qualitatively reshape the distribution. These results complement conditions previously derived for decoy-site binding (Bokes and Singh, 2015).

The outline of the paper is as follows. In Section 2, the stochastic process is constructed and an integro-differential master equation for its probability density function is formulated. In Section 3, we provide exact analytic results based on the analysis of the master equation. Section 4 builds on the analytical results to provide tractable approximations in the opposite scenarios of slow and fast sequestration kinetics. Section 5 presents the implications of these theoretical results and Section 6 concludes the paper.

## 2 Model formulation

We model bursty protein expression with sequestration by a hybrid, piecewise– deterministic process based on the schematic in Figure 1. Subsection 2.1 specifies the stochastic and deterministic parts of the process. Subsection 2.2 provides a deterministic fixed-point analysis. Subsection 2.3 derives the master equation for the time evolution of the protein probability density function (pdf).

**Figure 1.**
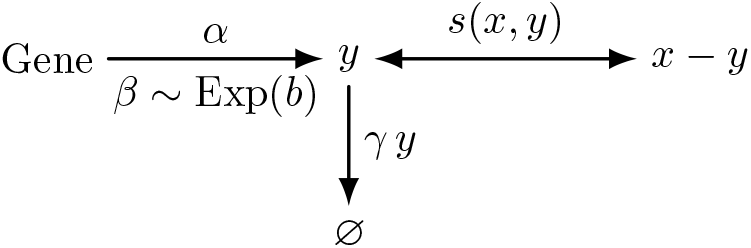
Model schematic: total protein *x* is partitioned into free (*y*) and sequestered (*x* − *y*) pools. Bursts arrive to *y* at rate *α* with sizes *β* ∼ Exp(*b*). Only free protein decays at rate *γ y*. Reversible sequestration transfers mass between *y* and *x* − *y* with net rate *s*(*x, y*). This can account for both non-cooperative and cooperative sequestration.

### 2.1 Gene expression process

We track three concentrations: *x*(*t*) for total protein, *y*(*t*) for free protein, and *z*(*t*) for sequestered protein, all nonnegative functions of time *t*. Since

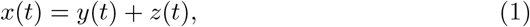

two variables suffice; we use the pair (*x*(*t*), *y*(*t*)).

Protein production is stochastic and occurs in random bursts. Bursts arrive as a Poisson process with rate (intensity) *α*:

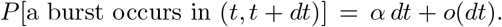

Burst sizes *β* are independent and exponentially distributed (Friedman et al., 2006)

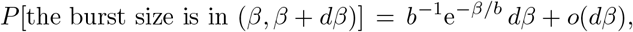

where *b* is the mean burst size. If a burst occurs at time *t*, the total and free concentrations increase by *β*:

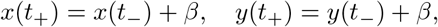

where *t*_−_ and *t*_+_ denote the left and right limits, respectively.

Between bursts, free proteins undergo decay and sequestration, modelled deterministically by

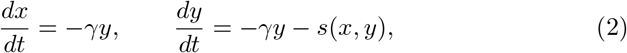

where *γ* is the decay rate constant. Only free proteins are degraded, while sequestered ones are assumed to be protected from decay. This assumption reflects situations in which sequestration provides physical or functional shielding from active degradation. In contrast, if protein loss were dominated by dilution (e.g., due to cell growth), both free and sequestered forms would experience comparable decay (Zhang et al., 2025).

The net sequestration rate is

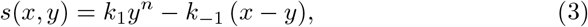

where *k*_1_ and *k*_−1_ are the sequestration and dissociation rate constants, and *n* ≥ 1 is the cooperativity. Subtracting the equations in (2) gives *dz/dt* = *s*(*x, y*) with *z* = *x* − *y*. Sequestration is linear when *n* = 1 and nonlinear when *n >* 1. The dimerisation case *n* = 2 covers, for example, formation of inactive ribosome complexes (Franken et al., 2017). The sequestration affinity is quantified by

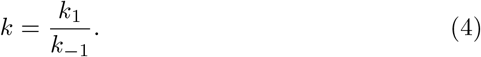

The same affinity *k* can arise from proportionally low or high rates. As illustrated in Figure 2, for large *k*_−1_ and *k*_1_ each burst is followed by a fast correction phase and then a slow decay.

**Figure 2.**
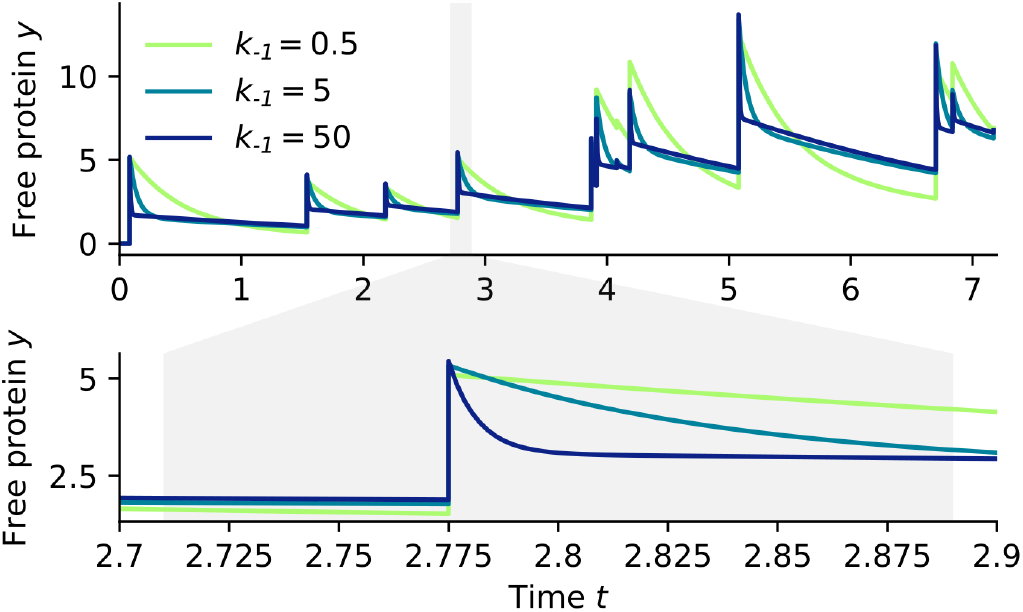
Sample trajectories of the free protein for different separations between sequestration kinetics and protein turnover. Under fast kinetics, sequestration is in quasi–steady state except briefly after bursts (one such interval is magnified in the lower panel). The inset reports *k*_−1_; the sequestration rate uses *k*_1_ = *k k*_−1_ with affinity *k* = 2. Other parameters: *n* = 1, *γ* = 1, *α* = 2, *b* = 3. Sample paths were generated with a hybrid stochastic simulation algorithm (Bokes et al., 2013).

### 2.2 Deterministic counterpart and fixed point analysis

Neglecting noise in the production process, we consider the deterministic counterpart of the stochastic model, where protein synthesis occurs continuously at its mean rate. The average production rate is *αb*, corresponding to the mean burst arrival rate *α* multiplied by the mean burst size *b*. The deterministic system is therefore

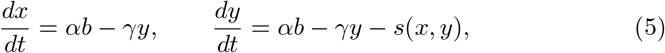

with *s*(*x, y*) given by (3).

A fixed point 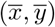 satisfies

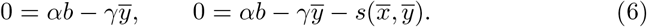

The first equation immediately gives

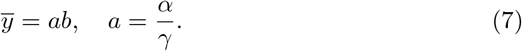

Thus, the mean free-protein level equals the product of the normalised burst frequency *a* and the mean burst size *b*, and it does not depend on *k*_1_, *k*_−1_, or *n*.

Substituting (7) into the second equation in (6) yields the steady-state constraint

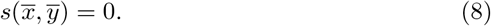

From (3), this implies

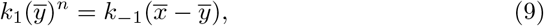

which defines a unique fixed point

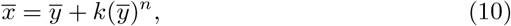

where *k* = *k*_1_*/k*_−1_ is the sequestration affinity.

To assess stability, we linearize (5) around 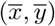. The Jacobian matrix is

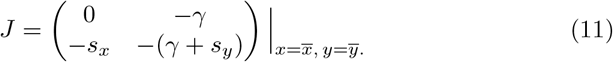

Here, *s*_*x*_ and *s*_*y*_ denote the partial derivatives of *s* with respect to *x* and *y*, respectively. Because *s*_*x*_ = −*k*_−1_ and 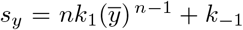, both eigenvalues of *J* are real and negative, so the fixed point is locally asymptotically stable for all positive rate constants, independently of the cooperativity exponent *n*.

Hence, the deterministic system possesses a unique, globally attracting steady state that represents the mass balance between production, degradation, and sequestration processes.

### 2.3 Master equation

The joint pdf *p*(*x, y, t*) for 0 *< y < x* and *t >* 0 of the stochastic process described in Subsection 2.1 satisfies the master equation (Zabaikina and Bokes, 2025)

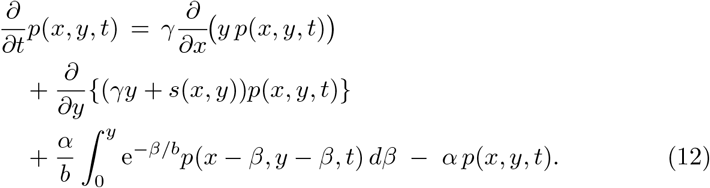

The right-hand side contains a deterministic (advective) part and a stochastic (integral) part. The deterministic term is the divergence of the pdf times the vector field in (2). The stochastic term is a convolution with the exponential burst-size density, applied along the diagonal in (*x, y*)-space (i.e., (*x, y*) ↦ (*x* − *β, y* − *β*)), because bursts increase *x* and *y* by the same amount (Zabaikina and Bokes, 2025).

At *t* = 0, the joint pdf is prescribed by

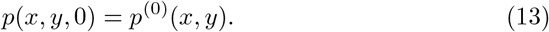

Typically, *p*^(0)^ is a delta distribution that places unit mass at deterministic initial concentrations.

A boundary condition is required for uniqueness. The boundary consists of states with no free protein (*y* = 0) or no sequestered protein (*y* = *x*). For an admissible solution of (12), the advective probability flux through the boundary must vanish, which gives

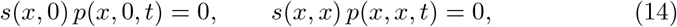

for any *x >* 0 and *t >* 0. Because *s*(*x*, 0) = − *k*_−1_*x <* 0 and *s*(*x, x*) = *k*_1_*x*^*n*^ *>* 0 by (3), the zero–flux conditions in (14) imply *p*(*x*, 0, *t*) = *p*(*x, x, t*) = 0; that is, the pdf vanishes on the boundary.

Parameters and variables used within this paper are listed in Table 1.

**Table 1.**
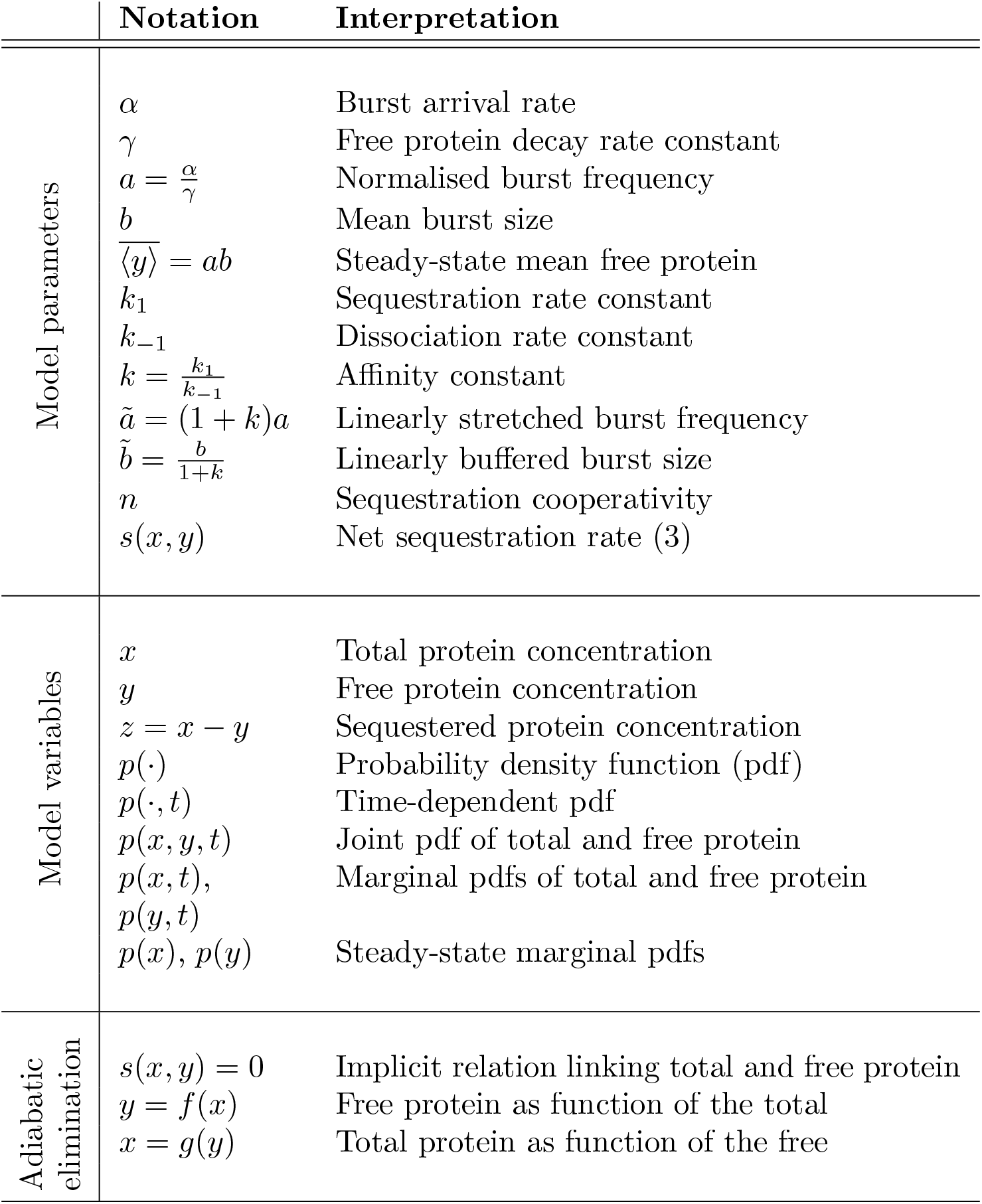
Parameters, variables, and quasi-steady-state relations.

|  | Notation | Interpretation |
| --- | --- | --- |
| Model parameters | $\alpha$ | Burst arrival rate |
| | $\gamma$ | Free protein decay rate constant |
| | $a = \frac{\alpha}{\gamma}$ | Normalised burst frequency |
| | $b$ | Mean burst size |
| | $\langle y \rangle = ab$ | Steady-state mean free protein |
| | $k_1$ | Sequestration rate constant |
| | $k_{-1}$ | Dissociation rate constant |
| | $k = \frac{k_1}{k_{-1}}$ | Affinity constant |
| | $\tilde{a} = (1 + k)a$ | Linearly stretched burst frequency |
| | $\tilde{b} = \frac{b}{1+k}$ | Linearly buffered burst size |
| | $n$ | Sequestration cooperativity |
| | $s(x, y)$ | Net sequestration rate (3) |
| Model variables | $x$ | Total protein concentration |
| | $y$ | Free protein concentration |
| | $z = x - y$ | Sequestered protein concentration |
| | $p(\cdot)$ | Probability density function (pdf) |
| | $p(\cdot, t)$ | Time-dependent pdf |
| | $p(x, y, t)$ | Joint pdf of total and free protein |
| | $p(x, t),$ | Marginal pdfs of total and free protein |
| | $p(y, t)$ | |
| | $p(x), p(y)$ | Steady-state marginal pdfs |
| Adiabatic elimination | $s(x, y) = 0$ | Implicit relation linking total and free protein |
| | $y = f(x)$ | Free protein as function of the total |
| | $x = g(y)$ | Total protein as function of the free |

## 3 Mean Invariance and Marginal Dynamics

We next analyze basic properties of the model. In Subsection 3.1, we show that the free-protein mean is independent of sequestration kinetics. In Subsection 3.2, we derive an exact but unclosed master equation for the marginal distribution of the total protein. In Subsection 3.3, we do the same for the free protein. Closure schemes for these marginal equations are deferred to Section 4.

### 3.1 Mean invariance

Consider an integrable function *f* = *f* (*x, y*) of the protein concentration (total and free) at a time *t >* 0 and its expectation

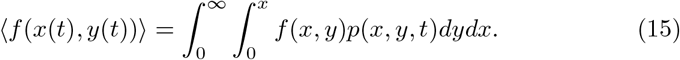

We use ⟨ · ⟩ for expectation and the conditioning notation ⟨ ·| ·⟩where needed. We multiply (12) by *f* (*x, y*) = *x* and integrate the product for all 0 *< y < x*, obtaining

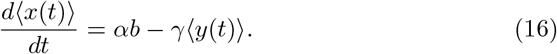

The boundary terms vanish by the zero–flux conditions in (14). For the steady-state (indicated by a bar) free protein mean we have:

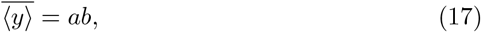

so that the stationary mean free protein agrees with the deterministic fixed point (7), which is independent of the sequestration parameters *k*_−1_, *k*_1_, and *n*.

### 3.2 Marginal total protein distribution

The marginal total protein pdf is defined by:

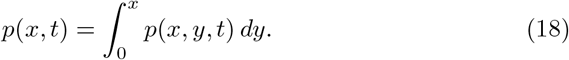

We use the same letter *p* for all pdfs, distinguishing them by their arguments. Integrating (12) over *y* ∈ (0, *x*) yields

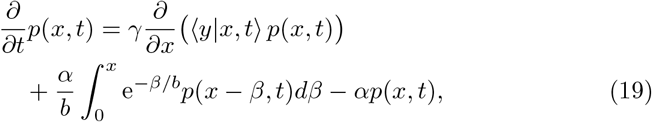

where the expected value of free protein *y* conditioned on the total protein *x* at time *t* is given by:

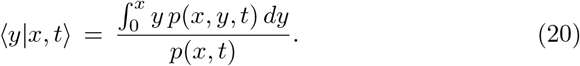

Equation (19) is exact, but not closed because the conditional expectation in (20) depends on the unknown joint pdf *p*(*x, y, t*).

### 3.3 Marginal free protein distribution

The marginal free protein pdf is given by

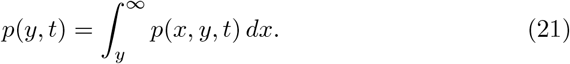

Integrating (12) over *x* ∈ (*y*, ∞) yields

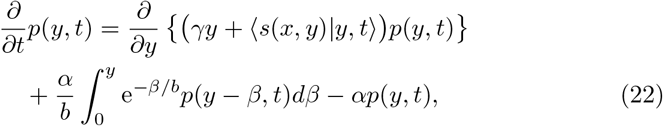

Where

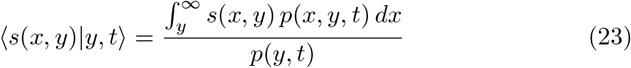

gives the conditional expectation of the net sequestration rate. As with the total-protein equation, (22) is exact but not closed because (23) refers to the joint pdf.

## 4 Limit Regimes and Adiabatic Reduction

Analytical progress is possible in two opposing limits of the sequestration kinetics. The simple case of *slow* sequestration (*k*_1_, *k*_−1_ → 0 at fixed affinity *k*) is treated in Subsection 4.1. The more interesting case of *fast* sequestration (*k*_1_, *k*_−1_ → ∞ at fixed *k*) is analysed in Subsection 4.2.

### 4.1 Slow sequestration

When *k*_1_ and *k*_−1_ are sufficiently small, sequestration contributes only higher– order corrections to (22). Neglecting the term (23) yields the closed equation

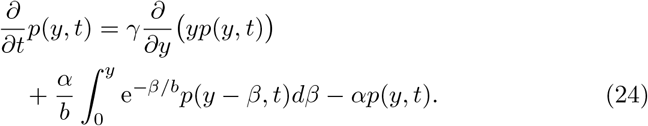

At steady state, (24) is solved by the gamma pdf with shape *a* and scale *b*:

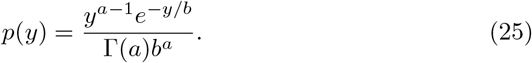

This is the well-known stationary distribution for bursty production with linear decay (Friedman et al., 2006), so slow sequestration leaves the steady-state free-protein distribution essentially unchanged.

The effective sequestration rate can be obtained by averaging over the above gamma distribution:

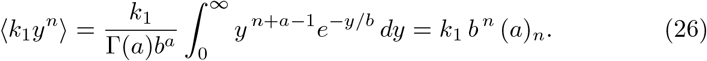

Here

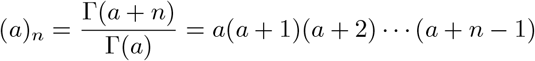

denotes the rising factorial (the Pochhammer symbol).

In the limit of very slow reaction rates, the dynamics of the sequestered protein become effectively deterministic:

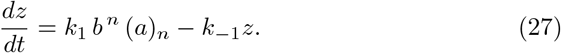

At steady state, the sequestered protein is thus deterministically fixed at

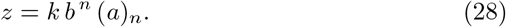

Interestingly, this expression deviates from the classical deterministic fixed point *z* = *k*(*ab*)^*n*^.

In the slow-sequestration regime, the piecewise deterministic model is expected to be inaccurate for the sequestered protein, since rare binding and unbinding events are treated as continuous processes. In a fully stochastic description, intrinsic binding–unbinding noise would instead give rise to a Poisson distribution with mean (28).

### 4.2 Fast sequestration

When *k*_1_ and *k*_−1_ are large, sequestration rapidly equilibrates. We adopt an adiabatic elimination in which the fast subsystem is at quasisteady state:

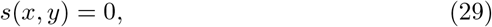

where the net sequestration rate on the left-hand side of (29) is defined by (3).

Using the implicit function theorem on (29), we express the free concentration as a function of the total:

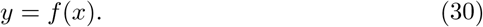

The function is generally non-elementary, as it is the inverse of

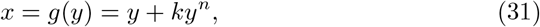

which gives the total protein as function of the free protein.

The adiabatic closure approximates the joint pdf by a delta mass along the slow manifold:

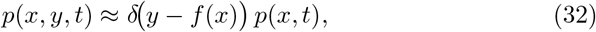

so that the delta distribution accounts for the functional relation (30). Substituting (32) into (20) gives

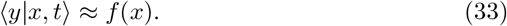

Substituting (33) into (19) gives the adiabatically reduced master equation

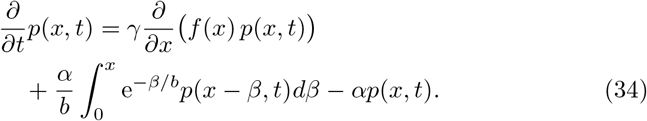

Its steady state has the form (Bokes and Singh, 2019)

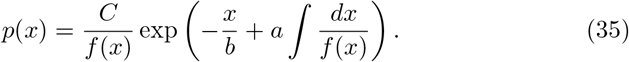

The free protein distribution is obtained by the transformation rule

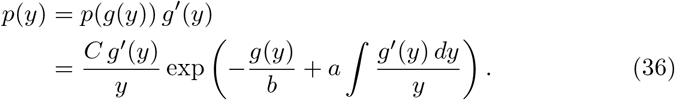

For *n* = 1, the transformation (31) is linear and we obtain a gamma distribution with modified parameters:

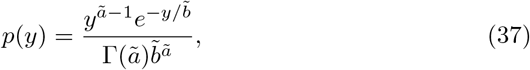

where

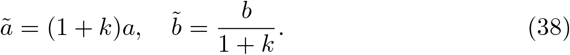

For *n >* 1, the transformation (31) is non-linear and we obtain a novel distribution:

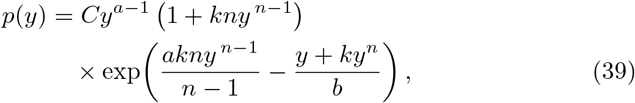

where *C* is determined by normalisation, 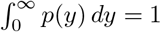.

## 5 Effects of Sequestration Kinetics and Cooperativity on Protein Distributions

We study the stationary distribution of the free protein *y* in a bursty expression model extended with sequestration. The mean is independent of sequestration kinetics and equals the product of burst frequency and mean burst size (Eq. 17).

Below we summarise how sequestration kinetics shapes other distributional properties. Kinetics are parameterised by the sequestration rate *k*_1_, the dissociation rate *k*_−1_, and the cooperativity coefficient *n* (Eq. 3). We treat the non–cooperative case *n* = 1 and the cooperative case *n >* 1 separately.

### 5.1 Non–cooperative sequestration (*n* = 1)

We consider the free–protein pdf *p*(*y*) = *p*(*y*; *k*_−1_) as a function of the dissociation rate *k*_−1_ while fixing the affinity *k* and setting *k*_1_ = *k k*_−1_. We examine the behaviour as *k*_−1_ varies from low (slow sequestration) to high (fast sequestration).

In the slow–sequestration limit, the distribution matches the no–sequestration case: a gamma law with shape *a* and scale *b* (Fig. 3, *k*_−1_ → 0). Thus, fast fluctuations are insensitive to slow sequestration.

**Figure 3.**
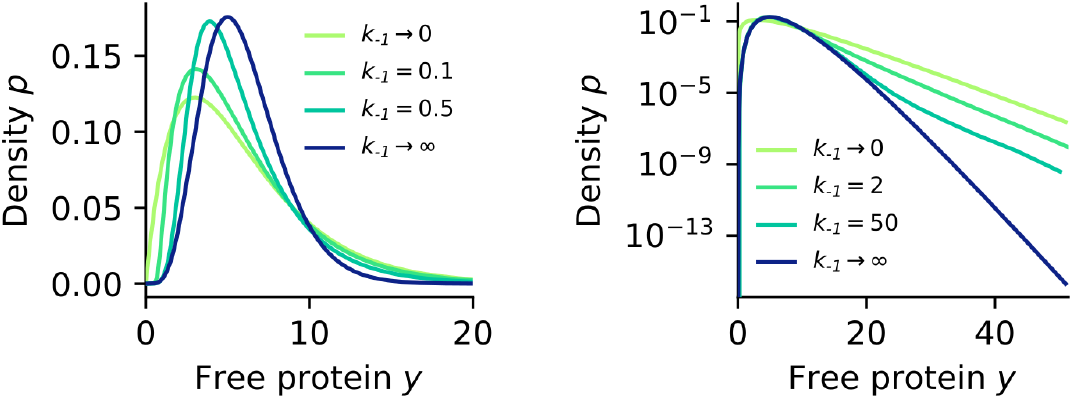
Steady–state distributions of the free protein with non–cooperative sequestration (*n* = 1) on linear (left) and logarithmic (right) scales. Faster kinetics drive the distribution from a high–noise to a lower–noise gamma form, while the right tail remains governed by the high–noise limit. The inset shows *k*_−1_; we set *k*_1_ = *k k*_−1_ with fixed *k* = 2. Other parameters: *γ* = 1, *α* = 2, *b* = 3. Distributions are histogram estimates from a hybrid stochastic simulation (Bokes et al., 2013) over total time *T*_max_ = 7 × 10^6^ with step Δ*t* = 0.005. The right panel uses “logarithmic binning” with geometrically increasing bin widths to reduce tail noise (Newman, 2004; Clauset et al., 2009).

In the fast–sequestration limit, the distribution is a lower–noise gamma (37) with shape 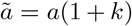 and scale 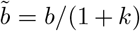. The parameter change reflects buffered burst sizes and an effective increase in burst frequency due to reduced decay of bound protein.

To characterise the transition from high–noise to low–noise gamma as kinetics speed up, we examine the logarithmic derivative of the pdf. On a log scale (Fig. 3, right), one observes a distinct slope in the *k*_−1_, *k*_1_ → ∞ limit and a different slope for finite rates (as *y* → ∞).

From the buffered gamma (37), the limiting logarithmic derivative is

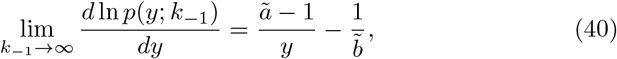

where 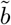 is defined by (38). For finite *k*_−1_ *>* 0, simulations indicate that as *y* → ∞ the logarithmic derivative approaches the unbuffered gamma slope

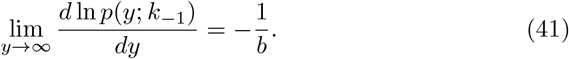

Letting *k*_−1_ → ∞ in (41) and *y* → ∞ in (40) yields a non-commutativity relation:

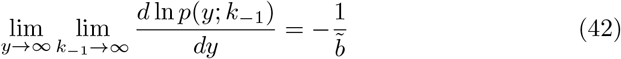

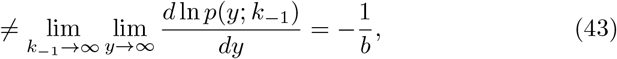

which highlights the nontrivial dependence of *p*(*y*) on *k*_−1_. In particular, simple families such as the gamma cannot capture this non–commutativity. Thus, although the limiting cases *k*_−1_, *k*_1_ → 0 and *k*_−1_, → ∞ *k*_1_ are gamma, for finite kinetics *p*(*y*) is novel and nontrivial.

### 5.2 Cooperative sequestration (*n >* 1)

We now compare cooperative to non–cooperative sequestration under fast kinetics, i.e. *k*_−1_, → ∞ *k*_1_ with fixed *k* = *k*_1_*/k*_−1_. In this regime, the cooperative case has the exact pdf (39), whereas the non–cooperative case yields the buffered gamma (37). To compare fairly, we match (*k, n*) pairs by equating the mean sequestered protein

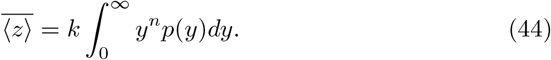

Because the free–protein mean (17) is kinetics–independent, matched pairs also share the same mean free and total protein; that is, they implement sequestration mechanisms with equivalent buffering capacity.

We examine two regimes: frequent bursts (burst frequency per lifetime *a >* 1) and rare bursts (*a <* 1). We visualise the distributions in Fig. 4 and report dispersion and skewness via

**Figure 4.**
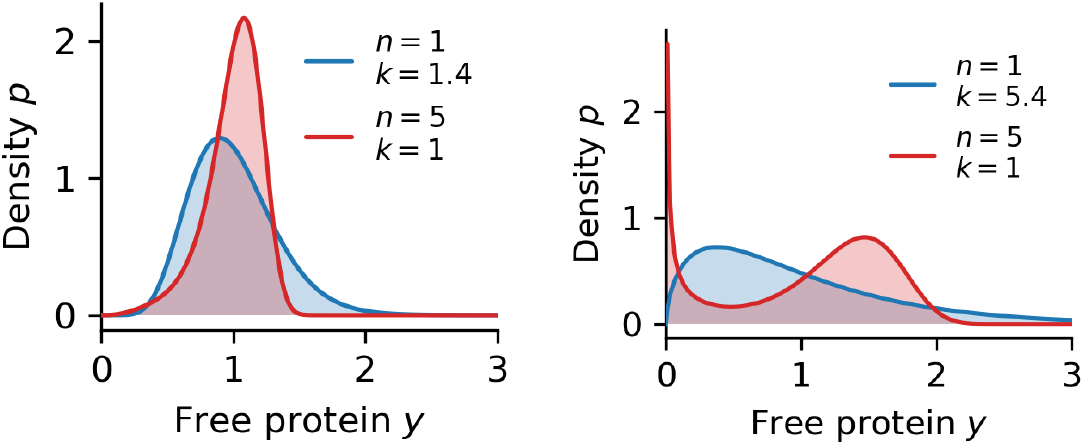
Steady–state free–protein distributions for fast non–cooperative (*n* = 1) and fast cooperative (*n >* 1) sequestration, with affinities chosen to match mean free, sequestered, and total protein. Cooperative sequestration reshapes the gamma law into left–skewed or multi–modal forms. Mean free protein ⟨ *y* ⟩= 1; burst size *b* = 1*/*4 (left) and *b* = 4 (right).

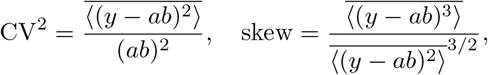

where 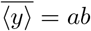 is the free protein mean. In the frequent–burst case (Fig. 4, left), increasing *n* reduces noise from CV^2^ = 0.1 at *n* = 1 to CV^2^ = 0.04 at *n* = 5, and flips the skewness from positive (skew = 0.65) to negative (skew = − 0.83). In the rare–burst case (Fig. 4, right), cooperative sequestration produces bimodality. As a result, noise reduction and skewness reversal are weaker (CV^2^ = 0.63, skew = 1.58 for *n* = 1 vs. CV^2^ = 0.4, skew = −0.39 for *n* = 5).

### 5.3 Bimodality parameter regions

To derive conditions for bimodality, we analyse the logarithmic derivative of the free–protein pdf as a function of the parameter vector 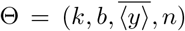. Differentiating (36) gives

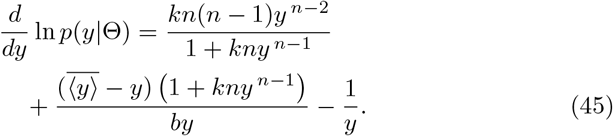

A unimodal–to–bimodal transition occurs at degenerate critical points satisfying

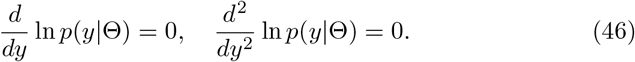

These two equations determine *y* and a three–dimensional hypersurface of critical points in the four–dimensional parameter space. Projecting to the (*k, b*)– plane gives a curve that separates unimodal and bimodal regions (Fig. 5).

**Figure 5.**
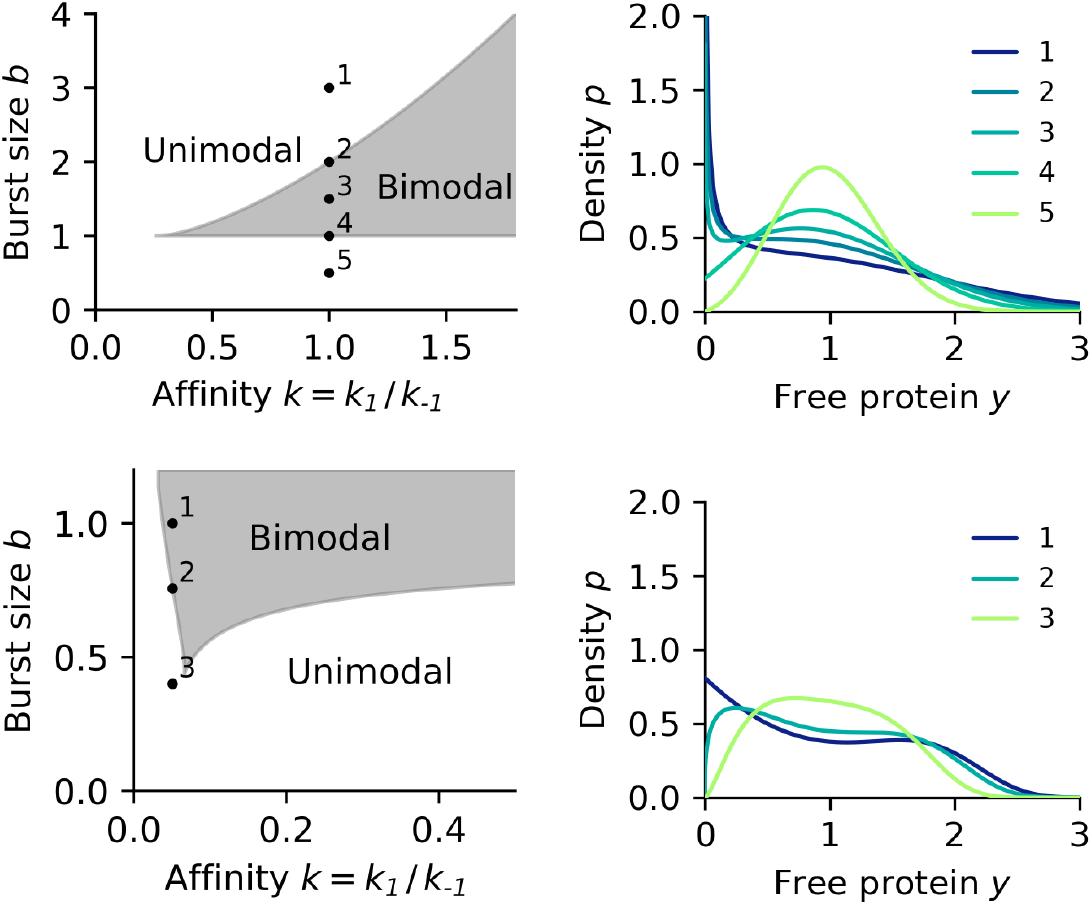
Left: unimodal and bimodal regions for *n* = 2 (top) and *n* = 5 (bottom). Right: densities corresponding to selected points in parameter space. The mean free–protein concentration is fixed at 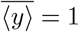.

Given that the deterministic model possesses a unique asymptotically stable fixed point, the bimodality exhibited by the stochastic model is noise-induced (Singh, 2012). Exploration of the parameter space indicates that the noise-induced bimodality emerges for large burst sizes (equivalently low burst frequencies). The extent of the bimodal region grows with cooperativity *n*. For dimerisation (*n* = 2), bimodality requires 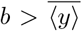 (i.e. *a <* 1) and sufficiently large affinity *k*. For higher–order multimers (*n* = 5), bimodality appears already at smaller *b* (higher burst frequency) and lower *k*.

## 6 Conclusion

We analyzed stochastic gene expression with sequestration and showed how kinetics and cooperativity shape the stationary distribution of free protein. The mean level is invariant to sequestration kinetics and equals the product of burst frequency and mean burst size. For non-cooperative sequestration, slow kinetics recover the classical gamma law, whereas fast kinetics yield a lower-noise gamma with a non-commuting tail limit. For cooperative sequestration, fast kinetics reshape the distribution, producing left skewness or bimodality, with parameter regions identified by degenerate critical points. These results clarify when sequestration buffers noise and when it induces qualitative changes in variability, offering testable signatures for inference from single-cell data.

